# *Wolbachia*-infected pharaoh ant colonies have higher egg production, metabolic rate, and worker survival

**DOI:** 10.1101/2023.01.31.526493

**Authors:** Rohini Singh, Sachin Suresh, Jennifer H. Fewell, Jon F. Harrison, Timothy A. Linksvayer

## Abstract

*Wolbachia* is a widespread endosymbiotic bacteria with diverse phenotypic effects on its insect hosts. *Wolbachia* also commonly infects social insects, where it faces unique challenges associated with its hosts’ caste-based reproductive division of labor and colony living. Here we dissect the benefits and costs of *Wolbachia* infection on life-history traits of pharaoh ants, *Monomorium pharaonis*. Pharaoh ants are relatively short-lived and show natural variation in *Wolbachia* infection between colonies, thereby making them an ideal model system for this study. We quantified effects on the lifespan of queen and worker castes, the egg-laying rate of queens across queen lifespan, and the metabolic rates of whole colonies and colony members. Newly-infected queens laid more eggs than uninfected queens but had similar metabolic rates and lifespans. Surprisingly, infected workers outlived uninfected workers. Infected colonies were more productive due to increased queen egg-laying rates and worker longevity, and infected colonies had higher metabolic rates during peak colony productivity. While some effects of infection, such as elevated colony-level metabolic rates may be detrimental in more stressful conditions, we did not find any costs of infection under laboratory conditions. Overall, our study emphasizes the beneficial effects of *Wolbachia* on colony-level growth and metabolism in this species.

## Background

*Wolbachia*, a widespread maternally-inherited endosymbiotic bacteria, is best known for its ability to manipulate a wide range of hosts [1,2]. *Wolbachia* is estimated to have infected more than 65% of all insect species but is also widespread in other invertebrates such as crustaceans, arachnids, and nematodes [3]. This bacterium often manipulates the host reproductive systems, thereby causing cytoplasmic incompatibility between infected and uninfected mates, killing or feminizing infected males, causing female-biased sex ratios, or inducing parthenogenesis[4,5]. *Wolbachia* can also have fitness-enhancing effects for the host, such as increased host fecundity and survival, conditional on the *Wolbachia* strain, host genotype, host species, and environment[6–12].

Even though *Wolbachia* infects an estimated one-third of all ant species[13],we have a limited understanding of the phenotypic effects of *Wolbachia* on ants and other social insects[13,14]. As in solitary insects, *Wolbachia* has often been considered to be a manipulator of reproductive strategies in multiple species of ants[15–18]. For instance, *Wolbachia*-infected species are more likely to have dependent colony foundation, where two or more queens start a nest site together, which is associated with changes in patterns of colony-level resource investment and queen phenotypes (Treanor and Hughes, 2019). Alternatively, as recently shown in the ghost ant, *Tapinoma melanocephalum, Wolbachia* may also be a nutritional symbiont ([19] and thus, may have beneficial effects in some ant species. Overall, when compared to solitary insects, the distinct biology of eusocial insects, specifically their reproductive division of labor and obligately cooperative lifestyle, may alter the types of *Wolbachia*-induced effects that occur in queens and workers in social insect colonies [13,15,16,18,20–25]Treanor & Hughes, 2019; [13,15,16,18,20–25].

In the current study, we dissect individual and colony-level benefits and costs of *Wolbachia* infection in one of the most well-studied invasive ants, *Monomorium pharaonis*. We previously showed that infected *M. pharaonis* colonies have queen-biased sex ratios [26,27] and higher colony growth and reproductive potential, with possibly increased reproductive senescence of infected queens [27]. In the current study, we quantified the effects of *Wolbachia* infection on the fecundity of queens across their lifespan, the metabolic rates of colonies and colony members at two different colony life cycle stages (i.e. in colonies with young versus mature queens), and the lifespan of queens and workers. Overall, we sought to determine if higher colony-level productivity of infected colonies may be explained by higher egg-laying of infected queens and if that has costs that may be specific to certain castes and colony life cycle stages.

## Methods

### *Source of infected and uninfected* Monomorium pharaonis *colonies and ant husbandry*

To initially produce a population of colonies with known *Wolbachia* infection status, where genetic background and infection status were relatively uncoupled, we systematically intercrossed colonies that were naturally infected or uninfected with *Wolbachia* for nine generations [27]. Next, we separately combined 15 of these colonies that were infected, and 14 of these colonies that were uninfected, to create two separate pools of ants that differed in *Wolbachia* infection but were genetically similar. We used these two pools to create replicate and genetically homogeneous source colonies of known infection status, which will be referred to as ‘source colonies’ from hereon (see Singh & Linksvayer 2020 for more information). We experimentally synchronized the age of the queens in these source colonies by removing all existing adult queens from the colonies to initiate production of new virgin queens and males and restart the colony life cycle. This produced queens of known and the same age across all the source colonies. These queen age-matched source colonies with known infection status were then used as sources for known-aged queens and to create replicate experimental colonies for all the assays performed below. All colonies used in the current study were reared at 27 ± 1 °C with approximately 50% relative humidity, and fed *ad libitum* synthetic agar diet (sugar:protein = 3:1; [28]) and dried mealworms (*Tenebrio molitor*) twice a week.

### Egg-laying by newly-mated queens

We first compared the numbers of eggs produced by newly-mated infected and uninfected queens over 50 days in replicate experimental colonies. These replicate experimental colonies were each created with 50 workers and 20 virgin queens that were mated with 15 virgin males. The number of eggs, larvae, pupae, and adults was censused in each colony when the queens were 5, 8, 11, 14, 17, 20, 23, 35, 39, 43, and 50 days old. We used a blind design for the study where the infection status of the experimental colony at the time of the census was unknown.

### Egg-laying by queens across their lifespan

We next compared egg-laying differences of 20 *Wolbachia*-infected and uninfected queens when the queens were one, three, four, six, and nine months old to quantify differences across the queens’ lifespan. We assayed total egg production over a 48-hour period by introducing 20 known-aged queens into replicate eggless experimental colonies. Each experimental colony was constructed with approximately 500 adult workers and 500 brood (larvae plus pupae), following a previously described protocol (Singh & Linksvayer 2020). After 48 hours, we censused the total number of eggs in these experimental colonies as a measure of queen egg-laying rate, and then returned the known-aged queens to their respective source colonies.

### Metabolic rate differences between infected and uninfected colonies and colony members

We compared metabolic rates of (a) infected versus uninfected whole colonies at two different colony life stages: colonies with one-month-old queens (n =12) and three-month-old queens (n = 8), representative of colonies with young and mature queens, respectively; and (b) different colony members, namely the brood and the queens from colonies with one-month-old queens. We estimated metabolic rates using flow-through respirometry [29] with a LiCor-7000 for whole colonies [30] and brood, and on LiCor-6252 for groups of 15 queens using the differential gas analyzer mode. We used dry CO_2_-free air at a flow rate of 125ml/min (25% of 500 ml/min flow controllers) for whole colonies and brood, and a flow rate of 50 ml/min (100% of 50 ml/min flow controllers) for groups of 15 queens. Additional detail on the respirometry is provided in the supplemental methods and in Fig. S1. We used source colonies to create replicate experimental colonies containing queens at the required ages (i.e. one month or three months old).

We quantified metabolic rates of intact whole colonies, just the brood from these colonies, and just the queens, at an early stage in the colony life cycle (i.e. when queens were one month old) as this is when growth curves are steepest (Singh and Linksvayer, 2020) and most likely to be affected by metabolic cost. We estimated the metabolic rates of 12 infected and 12 uninfected replicate experimental colonies with each colony having 20 one-month-old queens, approximately 250 workers, and 250 brood. We also estimated the metabolic rates of just the brood (i.e. eggs plus larvae plus pre-pupae plus pupae) from 11 infected and 11 uninfected colonies, after recording from the intact whole colony (i.e. also containing adult workers and queens).

We measured CO_2_ emission from one experimental colony per day and alternated between infected and uninfected experimental colonies to ensure that the queens were of similar age between the two groups at the time of measurement. We added a small water tube in the respirometer chamber along with the colony and the brood, to reduce any stress from possible dehydration for the brood. Finally, we also estimated metabolic rates of 14 *Wolbachia*-infected and 15 uninfected groups of 15 queens that were one-to two months old. We measured one to four groups of queens per day and alternated between infected and uninfected groups of queens to ensure even sampling across queen ages and colony life cycle stages.

We also estimated the metabolic rates of eight *Wolbachia*-infected and eight uninfected replicate experimental colonies with 20 three-month-old queens and approximately 500 adult workers and 500 brood (eggs to pupae) per colony. We recorded CO_2_ emissions from an infected and an uninfected colony per day. We chose this queen age since *Monomorium pharaonis* colonies peaked in their productivity and *Wolbachia*-infected colonies had increased reproductive investment than uninfected colonies [27]. Additional details can be found in the supplementary methods and Fig. S1.

### Effect of Wolbachia infection status on the survival of queens

We compared the survival of queens in 18 *Wolbachia*-infected and 16 uninfected small experimental colonies, each initiated with 20 queens that were each 2.5-months-old, as well as 50 workers. Once every three weeks, and then once a week after four months, we censused each colony and counted the number of eggs, larvae, pupae, and adults, in particular the number of surviving queens. At each census, we recorded the number of queens within each colony that survived (i.e. individual-level survival) and also whether *any* queens within each colony survived (i.e. group-level survival). We used a blind design for the study so that the infection status of each colony was unknown while collecting data.

### Effect of Wolbachia infection status on the survival of workers

To estimate the effect of *Wolbachia* infection status on worker survival, controlling for worker genotype (see Fig. S2), we compared the survival of 23 groups of 50 *Wolbachia*-infected workers and 25 groups of 50 uninfected workers. We set up these replicate worker groups using newly-eclosed adult workers and censused these worker groups once every three days, from August 30, 2019 to December 2, 2019. As for queen survival, at each census, we recorded the number of workers within each replicate group that survived (i.e. individual-level survival) and also whether *any* workers within each replicate group survived (i.e. group-level survival).

### Statistical analysis

We analyzed the data in R version 3.6.1 [31] with car [32] and lme4 [33] packages for regression analysis and ggplot2 [34] for visualization. We used survival [35] and survminer [36] packages to compare survival with log-rank tests and Cox proportional hazards, and visualize survival probabilities of experimental groups using Kaplan-Meir method.

A generalized linear mixed model framework (GLMM; [37] with Poisson error distribution was used to compare differences in egg-laying over time. For this model, the total number of eggs at each time point was used as the response variable, *Wolbachia* as a predictor variable, number of queens and age of the queens as fixed factors, and experimental colony ID as a random factor to account for repeated measures. To assess differences in egg-laying at specific time points, a generalized linear model framework (GLM; [37] with negative binomial error distribution was used with total number of eggs as response variable, *Wolbachia* as a predictor variable, and number of adult workers and queens as fixed factors.

We assessed the allometric relationship between metabolic rates of the whole colonies (microwatts) and mass of the colonies (grams) using a log-log plot (Fig. S3). Metabolic rates were estimated, in microwatts and microwatts per gram of the experimental group, from CO2 levels measured in ppm by assuming an oxyjoule of 19.87 J ml^-1^ O2 (respiratory quotient of 0.75) and standardized to 25°C assuming a Q_10_ = 2.0 [29]. We used a linear model framework (LM) to test the effects of *Wolbachia* infection, queen age, colony-level activity, colony mass, and colony size on estimates of metabolic rates. We computed the test statistic of individual factors in the linear model via ANOVA from the car package [32]. The dataset used in the study, the detailed R script for data analysis, and the output from the regression models can be found as supplementary information.

## Results

### Wolbachia-infected pharaoh ant queens lay more eggs early in their life cycle

Newly-mated *Wolbachia*-infected queens produced more eggs over time than uninfected queens (GLMER: *χ^2^* = 7.6, *p* = 0.005; Fig 1a). Specifically, *Wolbachia*-infected groups of queens had more eggs at queen ages of: 8-days-old (GLM: *χ*^2^= 4.42, *p* = 0.035), 23-days-old (GLM: *χ*^2^= 6.82, *p* = 0.009), 35-days-old (GLM: *χ*^2^= 5.57, *p* = 0.018), and 50-days-old (GLM: *χ*^2^= 4.81, *p* = 0.028). However, such egg-laying differences were observed only during the early life stage of the queens (Fig. 1b). Colonies with 1-month-old *Wolbachia*-infected queens produced more eggs compared to their uninfected counterparts (GLM: *χ*^2^= 5.88, *p* = 0.015), while experimental colonies with older queens, did not show significant differences (Fig. 1b).

**Fig 1.**
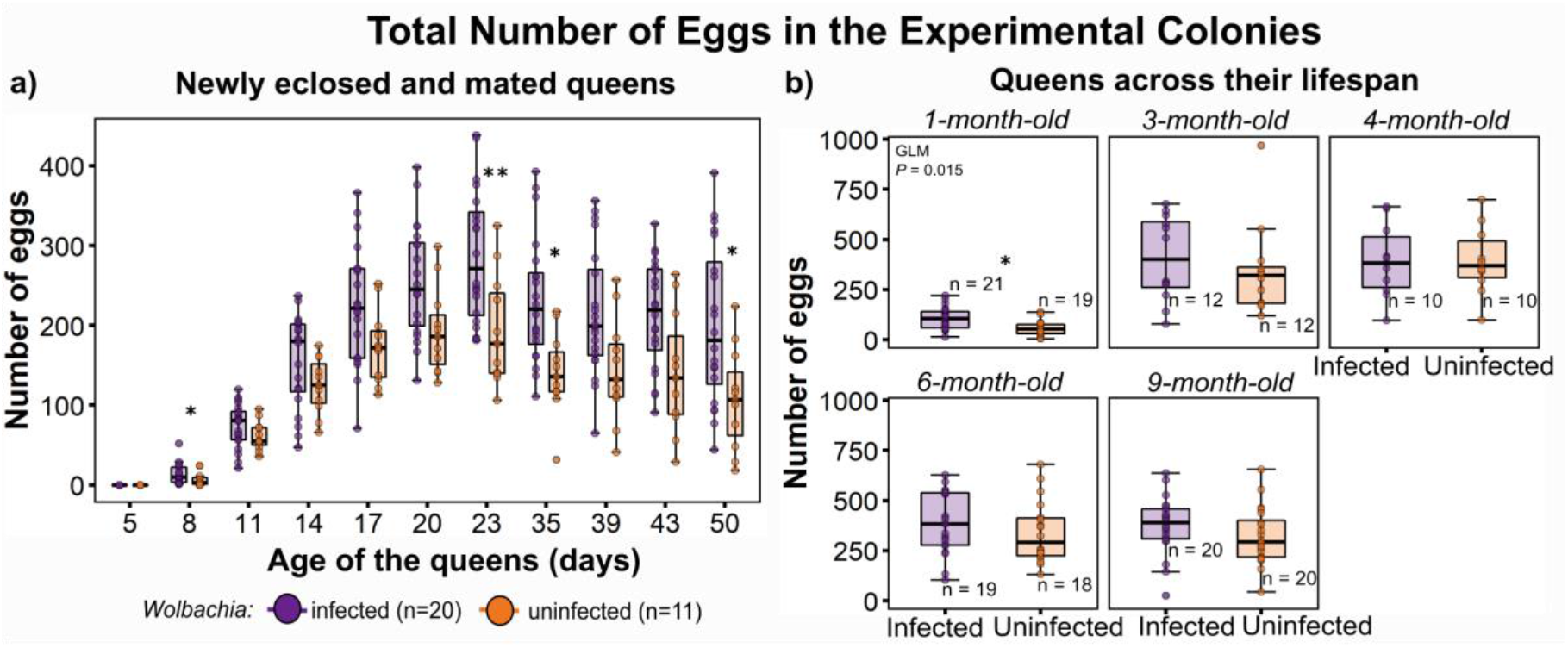
*Wolbachia*-infected queens lay more eggs soon after mating. (a) Groups of 20 newly-mated *Wolbachia*-infected queens lay more eggs than uninfected queens. (b) Colonies with 20 one-month-old *Wolbachia*-infected queens laid more eggs after 48 hours of adding the queens to the colonies. However, such differences were not observed when the queens were older. Box plot represents the quartile distribution of the raw data, the filled dots represent the individual raw values. For (a) *Wolbachia* color legend, along with the sample size (n) is included at the bottom of the graph. The x-axis represents the age of the queens in days and the y-axis represents the total counts of eggs in the colonies. For (b) the x-axis represents the *Wolbachia* infection status of the experimental colonies and the y-axis represents the total counts of eggs after 48 hours of adding the queens to the experimental colonies. Sample sizes (n) have been included on individual graphs. Significant differences due to *Wolbachia* infection, as computed from a GLM model, with *P* < 0.05 is represented by ‘*’ and with *P* < 0.01 is represented by ‘**’ on the graphs in (a) and (b).

### Wolbachia-*infected colonies have higher metabolic rates depending on the stage of the colony life cycle*

Metabolic rates (microwatts) of whole colonies showed hypometric scaling with mass (Fig. S3a) and had a scaling coefficient of 0.58 (95% CI, 0.45 - 0.71), which is within the expected range [38,39]. This means that the mass-specific metabolic rate (microwatts per gram) will decrease with increasing mass of the ant colony. In contrast, t, the scaling coefficient of metabolic rates (microwatts) of only the brood was 1.1 (95% CI, 0.32 - 1.94), which suggested that as brood mass increases, mass-specific metabolic rates will increase similarly to expectations by isometric scaling (Fig. S3b). Interestingly for the groups of queens, mass-specific metabolic rates did not show a significant scaling effect with mass, perhaps due to the small size variation in groups of 15 queens (Fig. S3c). For the remainder of the discussion, metabolic rates (microwatts) of colonies and different groups are discussed.

*Wolbachia*-infected pharaoh ant colonies with young queens (1- to 2-month-old) had similar metabolic rates as the uninfected colonies (LM: F = 0.57, *p* > 0.05; Fig. 2a). In contrast, *Wolbachia*-infected colonies with older queens (3-month-old) had higher metabolic rates than uninfected colonies (LM: F= 15.6, *p* = 0.002; Fig. 2b).

**Fig 2.**
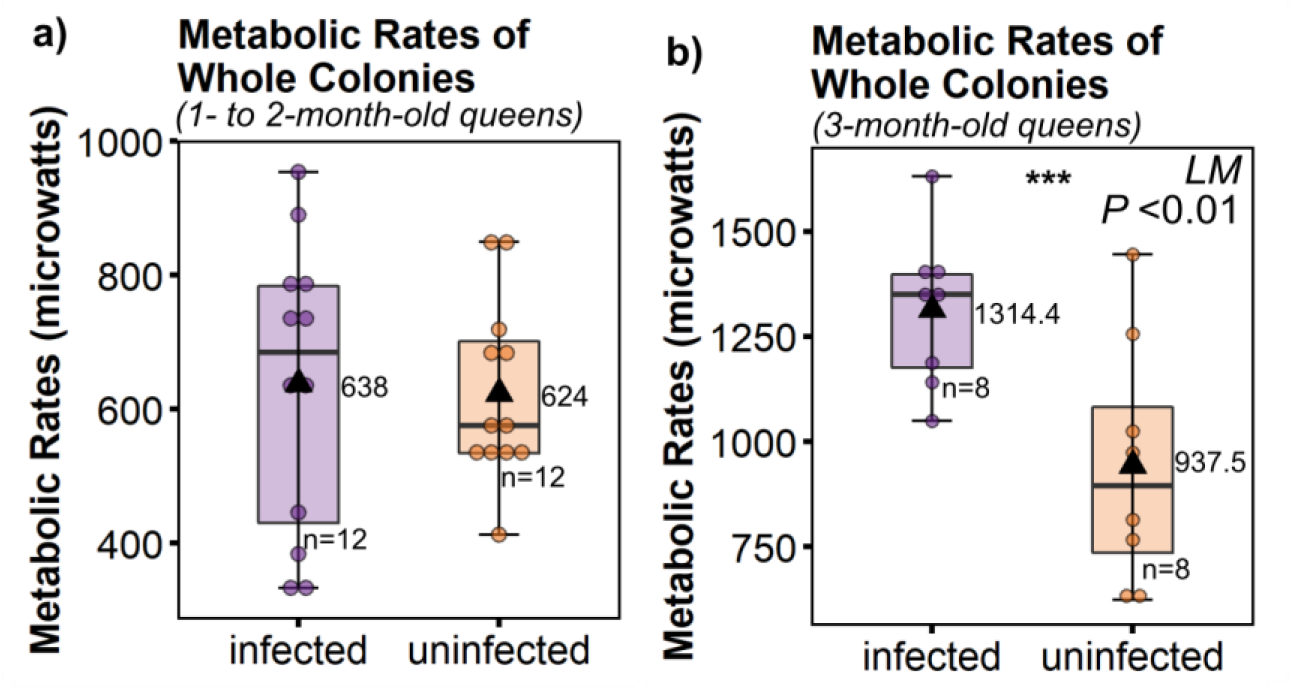
Metabolic rates (microwatts) differ between infected and uninfected groups but are dependent on colony life cycle stage and colony component. (a) Infected and uninfected whole colonies with 1- to 2-month-old queens have similar metabolic rates. (b) Infected colonies with 3-month-old-queens have higher metabolic rates than uninfected colonies. X-axis represents the *Wolbachia* infection status of the experimental group. Y-axis represents the metabolic rates of the groups in microwatts. Box plot represents the quartile distribution of the raw data, the filled dots represent the individual raw values. The filled black triangle in the box plot represents the mean, which is also numerically listed besides the box plot. ‘n’ represents the sample size for the accompanied box plot. ‘***’ represents the significant difference between infected and uninfected groups, as determined by a linear model, with *P* < 0.001.

We also compared the metabolic rates of different colony members when the colony was in early life cycle stages (1- to 2-month-old queens). Metabolic rates of the brood (eggs to pupae) increased with the age of queens initially present in the colonies (LM: F= 9.81, *p* = 0.006). Total brood mass also positively influenced brood metabolic rate (LM: F= 7.22, *p* = 0.016), and total number of brood had a marginal effect (LM: F= 3.22, *p* = 0.091). The metabolic rates of groups of queens increased with the age of the queens (LM: F= 16.63, *p* < 0.001) after statistically accounting for variation in mass of the queens. However, brood from these colonies did not show differences in metabolic rates when compared to uninfected brood (LM: F= 0.34, *p* > 0.05; Fig. S4a). *Wolbachia*-infected groups of 15 queens that were 1- to 2-month-old had similar metabolic rates as the uninfected queens (LM: F= 1.9, *p* = 0.18; Fig. S4b) with no significant interaction of queen age with *Wolbachia* infection (LM: F= 0.98, *p* > 0.05).

### Caste-specific survival differences due to Wolbachia

Despite differences in egg-laying of queens and in colony-level metabolic rates at some queen ages, *Wolbachia*-infected and uninfected queens had similar group- (Log-rank test, *p* = 0.8; Fig. 3a) and individual-level survival rates (GLMM, *χ*^2^ = 0.2, *p* <0.05; Fig. 3b). The estimated median survival of groups was 230 days for *Wolbachia*-infected queens and 206 days for uninfected queens (Fig. 3a). Within groups, the proportion of living queens over time was also similar between infected and uninfected groups (GLMM, *χ*^2^ = 0.2, *p* <0.05; Fig. 3b).

**Figure 3:**
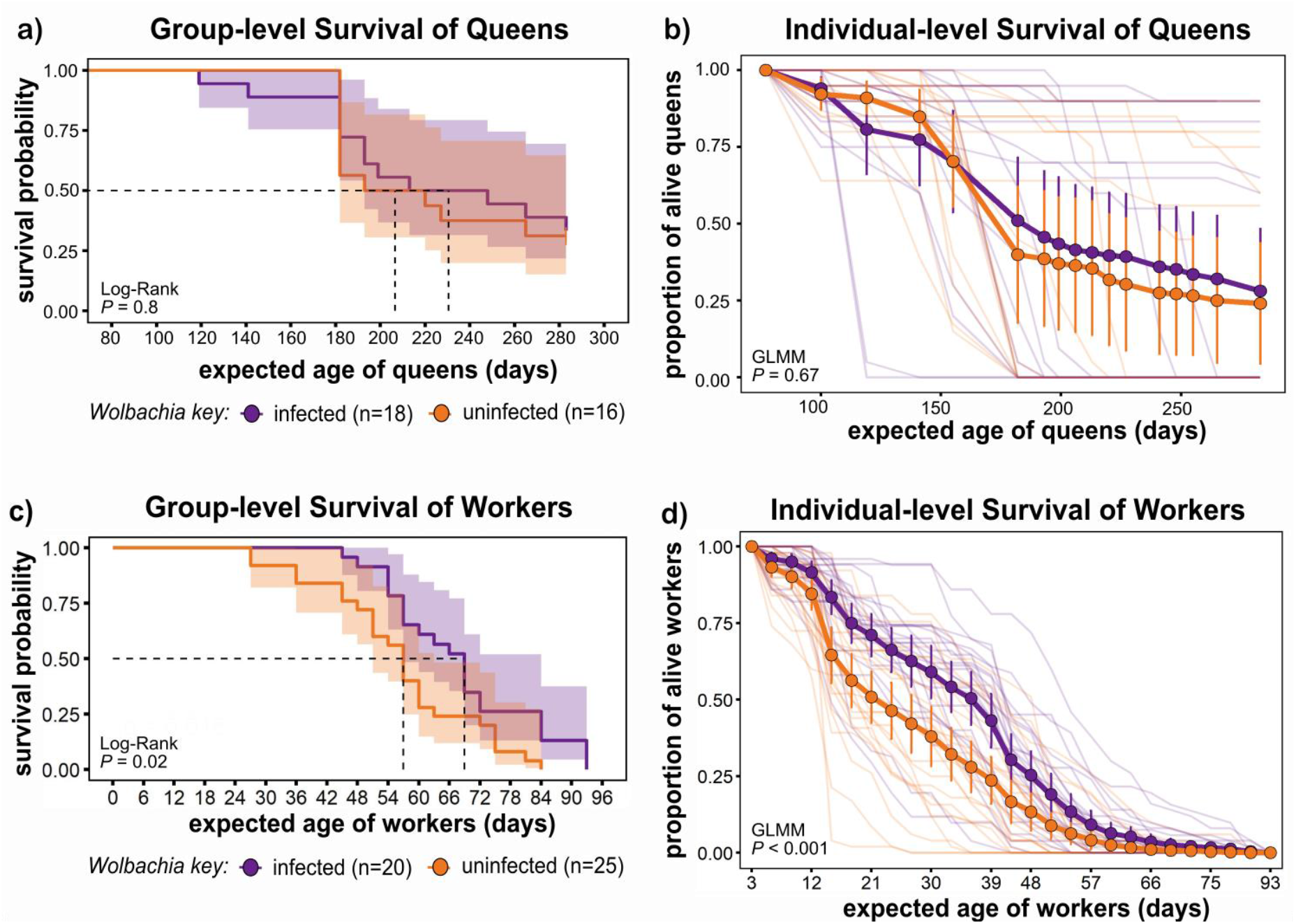
Survival differences are dependent on *Wolbachia* infection and caste. (a) Groups of infected and uninfected queens have similar survival probability. (b) Within the groups of queens, infected and uninfected groups have similar proportions of alive queens over time. (c) Infected groups of workers have a higher survival probability than uninfected groups of workers. (d) Within the groups of workers, infected groups have higher proportions of alive workers over time. X-axis represents the estimated age of queens (a, b) or workers (c, d). Y-axis represents the survival probability as estimated by Kaplan-Meier method (a, c) or proportion of alive queens (b) or workers (d). For (a) and (c) solid line represents the mean along with the 95% confidence interval (shaded area). The *P*-value using log-rank test with cox-proportional hazards model is listed on the bottom left corner of the graph. For (b) and (d), filled circles represent the mean value with 95% confidence interval (error bars). Solid dark line represents the mean trend and lighter lines represent the trend of individual groups. *P*-value estimate from GLMM is listed at the bottom left corner.

Infected workers had higher group-level survival (Log-rank test, *p* = 0.02; Fig 3c) and individual-level survival (GLMM, *χ*^2^ = 12, *p* < 0.001; Fig. 3d) than uninfected workers (Fig. 3c,d). The estimated median survival of groups was 69 days for infected workers and 57 days for uninfected workers. Groups of 50 *Wolbachia*-infected workers had a higher estimated survival probability than their uninfected counterparts. Within the group, a higher proportion of infected workers survived over time (GLMM, *χ*^2^ = 12, *p* < 0.001; Fig. 3d).

## Discussion

We compared individual- and colony-level life history traits of infected and uninfected *Monomorium pharaonis* colonies to elucidate the benefits and costs of *Wolbachia* infection. Newly-mated *Wolbachia*-infected queens produce more eggs without a metabolic cost. However, at a later colony life cycle stage (three-month-old queens), when colonies peak in their productivity and reproductive investment [27], infected colonies have higher metabolic rates. Despite increased egg-laying by queens and higher colony-level metabolic rates, *Wolbachia* infection was not associated with decreased queen lifespan. Interestingly, in workers, which are obligately sterile, *Wolbachia* infection was associated with longer lifespan. Thus, increased egg-laying rates by queens and longer worker lifespans contribute to the higher growth rate and productivity that characterizes *Wolbachia*-infected colonies [27].

Increased egg production by *Wolbachia*-infected queens may be caused by individual-level differences in the queens, such as increased stem cell differentiation or oogenesis, as has been shown in *Drosophila mauritiana* [40] and *Asobara tabida* [41], and/or differences in the ability of infected workers to rear more eggs. Cross-fostering infected queens with uninfected workers and *vice-versa* will be useful to tease apart the role of queens, workers, and queen-worker interaction on *Wolbachia*-induced phenotypes.

Given the increased egg-laying by infected queens and increased growth of infected colonies [27], we expected that infected colonies would have higher metabolic rates. Furthermore, we expected that this energetic cost would be exacerbated by the maintenance cost of *Wolbachia* [6,12,42]. We did find this pattern in the three-month-old colonies. However, we did not find differences in metabolic rates of infected and uninfected whole colonies, brood, and queens when the queens were young (one to two months old). This suggests that *Wolbachia* infection does not have detectable energetic costs, or perhaps *Wolbachia* offsets any costs through a nutritional symbiosis, as found in the bed bug (*Cimex lectularius;* [43,44], fruit fly (*Drosophila melanogaster*; [45], and the ghost ant (*Tapinoma melanocephalum*; [19]. Future studies comparing the metabolic rates of colonies and colony members across multiple colony life cycle stages will be helpful to better understand the energetic costs of *Wolbachia* infection.

A trade-off between fecundity and longevity has widely been observed within and between species, often assumed to be due to the costs of reproduction [46,47]. In contrast, social insects show a reversal in this fecundity-longevity tradeoff, where queens have high fecundity and long lifespans and workers are facultatively or obligately sterile and have short lifespans [48]. However, while we found an effect of *Wolbachia* infection on queen fecundity, we found no effect on queen lifespan. Surprisingly, we found that *Wolbachia*-infected workers, which are obligately sterile, did have longer lifespans. While both positive and negative effects of *Wolbachia* infection on the lifespan of solitary hosts have been observed [6,12,49], the mechanisms remain largely unknown. Since workers are obligately sterile in *M. pharaonis* colonies, the presence of infected fecund queens that live as long as uninfected queens may be beneficial for *Wolbachia*, as more infected individuals can be produced over time. Future studies teasing apart the mechanisms by which *Wolbachia* infection causes higher egg-laying rates in *M. pharaonis* queens with no effect on lifespan and longer lifespans in sterile *M. pharaonis* workers can lead to general insight into mechanisms linking reproduction, aging, and metabolism.

## Conclusion

We report that *Wolbachia* infection is beneficial for *Monomorium pharaonis* ant colonies, at least in relatively benign laboratory conditions, as infected young queens produced more eggs, infected colonies had higher metabolic rates during periods of peak productivity, and infected queens lived as long as the uninfected queens, while infected workers outlived uninfected workers. These phenotypic effects of infection suggest that *Wolbachia* may have adapted to exploit the ant reproductive caste system without exacting a detectable cost on its ant host. The phenotypic and fitness consequences of *Wolbachia* infection that we observed, if also observed under more natural and in particular more stressful conditions, are expected to be associated with the rapid spread of *Wolbachia* infection. However, the fact that *Wolbachia* infection is not universal in *M. pharaonis* colonies (Schmidt 2010) indicates that at least under some environmental conditions, there are presumably costs of infection that limit the spread of *Wolbachia* within *M. pharaonis* populations. Future experiments assessing the benefits and costs of *Wolbachia* under a variety of environmental conditions, especially stressful conditions, are needed to clarify these issues.

## Data accessibility

Data files and detailed R scripts can be accessed at Dryad

(https://datadryad.org/stash/share/4PxUOLVYZX-AtdvshO9tbqXqIOJMFWBxWkeMrkyR9h8).

## Author’s contributions

RS, JHF, JFH and TAL conceived and designed the study. RS collected the data. RS and SS analysed the data. RS, SS, and TAL drafted the manuscript, and RS, SS, JHF, JFH and TAL critically revised the manuscript.

## Acknowledgments

We are grateful to Dr. Juergen Liebig for letting us safely house the ants at Arizona State University, Xuaohui Guo in the Fewell lab and Trevor Fox in the Harrison lab at the Arizona State University for assisting with metabolic rate measurement, and the Linksvayer lab members for their feedback at various steps of the study.

