## Supplemental file 1 for "*Wolbachia*-infected pharaoh ant colonies have higher egg production, metabolic rate, and worker survival"

### Supplementary Figures

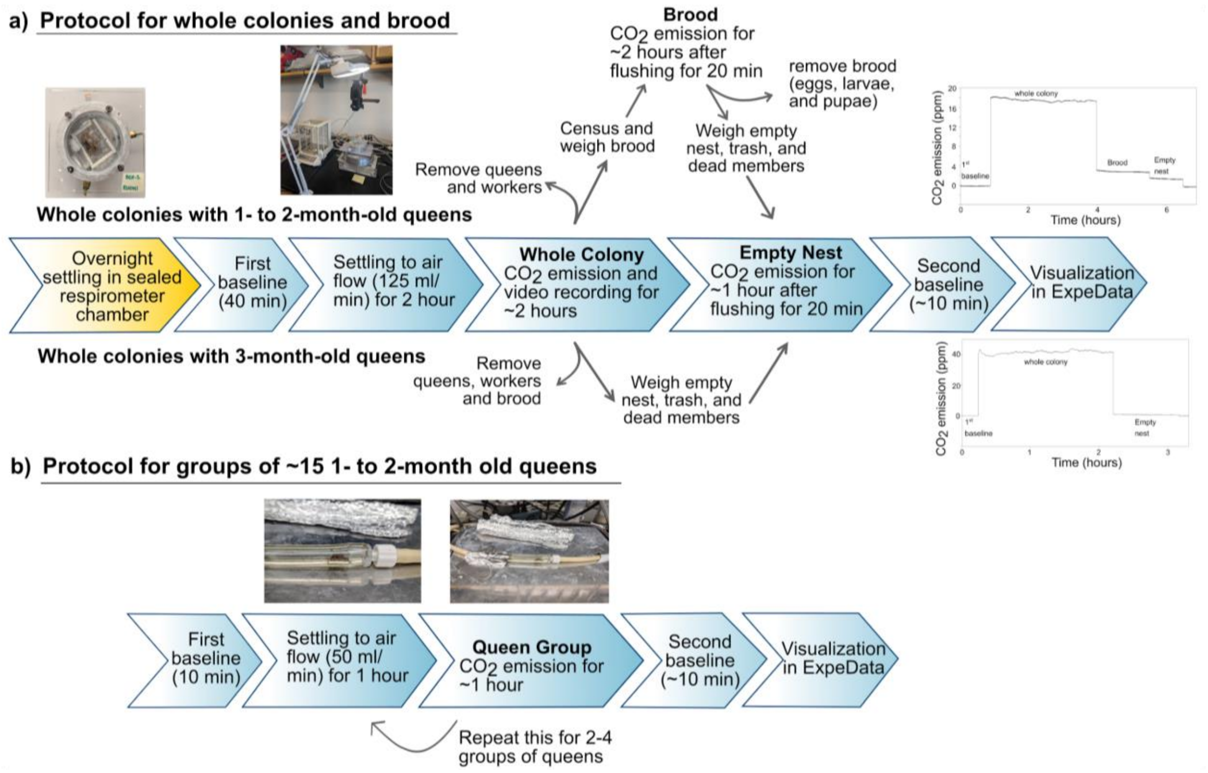


**Figure S1: Setup used for estimating metabolic rates.** (a) Detailed steps for measuring the $CO_2$ emission from whole colonies and brood with 1- to 2-month-old queens (top half) and whole colonies with 3-month-old queens (bottom half). (b) Detailed steps to measure $CO_2$ emission from groups of 1- to 2-month-old queens. Yellow color highlights the steps done a day prior to the measurement, whereas the blue color highlights the steps performed on the day of recording $CO_2$ emission on a respirometer.

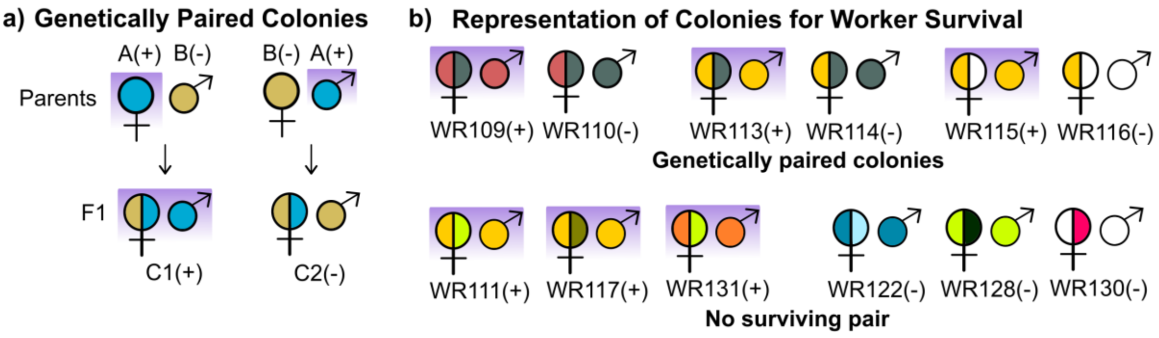


**Figure S2: Reciprocal crossing scheme to produce genetically paired *Monomorium pharaonis* colonies that differ in *Wolbachia* infection for comparing worker survival.** (a) We used a reciprocal crossing scheme to control for genotype when comparing *Wolbachia*-driven differences in life history traits of colonies and colony members. 'A' and 'B' represent sample parent colony ID of differing genotypes and 'C1' and 'C2' represent sample F1 colony ID. (b) A graphical representation of genetic diversity of the colonies used for comparing worker survival. We used 3 pairs of colonies that were expected to be genetically similar but have different *Wolbachia* infection status (top half). We also used colonies that did not have a surviving counterpart (bottom half). Each color represents a unique colony ID from heterogeneous stock colonies used for setting up reciprocal crosses. '(+)' following the colony ID and with a purple background means that the colony is infected with *Wolbachia*. '(-)' with a white background means that the colony is uninfected.

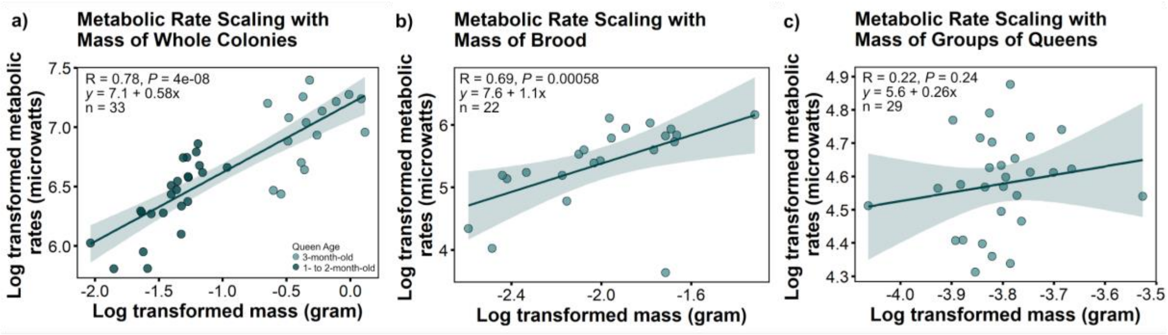


**Figure S3: Metabolic rate scaling with mass of the experimental group.** Log-log plot of metabolic rate with mass of (a) whole colonies with all our data combined (colonies with 1- to 3-month-old queens), (b) only the brood (from colonies with 1- to 2-month-old queens), and (c) groups of approximately 15 queens (1- to 2-month-old). '*R*' represents the Spearman Rank Correlation coefficient, '*P*' represents the significance of correlation, and 'n' represents the sample size. The regression line equation is represented on the top left corner in the format of 'y = x + mc', where 'm' is the scaling coefficient.

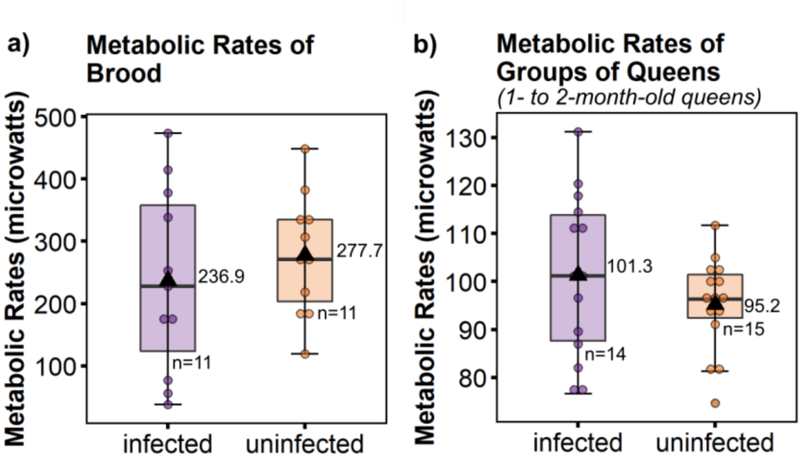


**Fig S4. Metabolic rates (microwatts) did not differ between infected and uninfected groups early in the colony life cycle.** (a) Similar metabolic rates of brood from the colonies with 1- to 2-month-old queens. (b) Similar metabolic rates of groups of 15 1- to 2-month-old queens. X-axis represents the *Wolbachia* infection status of the experimental group. Y-axis represents the metabolic rates of the groups in microwatts. Box plot represents the quartile distribution of the raw data, the filled dots represent the individual raw values. The filled black triangle in the box plot represents the mean, which is also numerically listed besides the box plot. ‘n’ represents the sample size for the accompanied box plot.

### Supplementary methods

*Comparing metabolic rate differences*

We performed respirometry measurements of whole colonies and brood in Dr. Jennifer Fewell's Lab and respirometry measurements of groups of queens in Dr. Jon Harrison's lab at Arizona State University (Tempe, AZ) by transporting experimental *Monomorium pharaonis* colonies in 50 ml falcon tubes in a cabin bag on a flight. These colonies were stored in Dr. Juergen Leibig's ant room chamber at Arizona State University (Tempe, AZ) at 26$^{o}$C with ± 60% relative humidity. The colonies were fed *ad libitum* synthetic agar along with dried mealworms twice a week. We transported the colonies twice, once in February 2018 when the colonies had 3-month-old queens and then again in November 2019 when the colonies had 1-month-old queens.

A day before recording $CO_2$ emissions from whole colonies, we censused and weighed the whole colony and then kept them overnight in a sealed respirometer chamber to settle in the new space (Fig. S1). The inner wall of the respirometer chamber was coated with a layer of fluon and the internal openings of the chamber were covered by a fine mesh to prevent ant escapes. For colonies with young queens (<2-month-old), since we wanted to record $CO_2$ emissions from the brood after recording from an experimental colony, we also transferred a small water tube into the respirometer chamber for this overnight settling and $CO_2$ emission recording the next day to alleviate any possible stress due to long periods of absence of humidity (Fig S1a). We started the $CO_2$ emission recording by first measuring a 40-min long first baseline on LiCor-7000, which served as a reference for any differences in $CO_2$ levels between the reference and sample cells of LiCor-7000 (Fig S1a). Following this we connected the respirometer chamber with our experimental colonies to the LiCor-7000 setup and allowed the colonies to settle for another 2 hours at a flow rate of 125 ml/min. After this we started recording the $CO_2$ emission from the respirometer chamber, ambient temperature, and the humidity within the respirometer chamber for approximately two hours till the colonies showed a stable emission. Post this, the colonies were taken out from the respirometer chamber and we used two different approaches between the colonies with queens younger than 2 months and colonies with 3-month-old queens. For colonies with 3-month-old queens, we returned the whole colony back to its original box, and weighed the empty nest, with any possible trash and dead colony members before placing the empty nest back to the respirometer chamber (Fig S1a). For colonies with queens younger than 2 months, we moved all the queens and workers from the colony back to its original box, censused and weighed the brood (eggs, larvae, pre-pupae, and pupae) and then proceeded to record $CO_2$ emission from just the brood for approximately two hours in the same way as described above (Fig S1a). After recording from the brood, we returned all the brood back to its original colony box but retained any dead colony members and trash within the respirometer chamber and nest. For both our approaches, we weighed the empty nest with dead colony members and trash and then sealed them in the respirometer chamber. We then passed $CO_2$-free dry air at a flow rate of 125 ml/min for 20 min to flush any environmental $CO_2$ that made its way into the chamber during the entire process (Fig S1a). We recorded $CO_2$ emissions from the empty nest for approximately an hour as a reference for background $CO_2$ emissions (Fig S1a). After this we removed the respirometer chamber, connected the tubings to each other and recorded the second baseline for 10 min to account for any drift over the course of recording. We also recorded videos of the whole colonies over the course of $CO_2$ emission measurements to later compare activity level differences.

We also estimated metabolic rates of groups of approximately 15 young queens in 2019 by storing them in tubing with fine mesh on both ends to prevent ant escapes (Fig S1b). We recorded $CO_2$ emission rate from these groups of queens using the differential mode in LiCor-6252 gas analyzer by flowing dry $CO_2$-free air at 50 ml/min through mass flow controllers (50 ml/min max, set to 100%). The queens were collected from replicate experimental colonies which had already been measured and the queens were weighed before recording their $CO_2$ emission. We first recorded a 10-min long first baseline. Following this, we connected the tube with the group of queens, allowed the group to settle to the airflow for about an hour and then started recording $CO_2$ emission from this group for another hour before returning them back to their respective colonies (Fig S1b). After this we added another group of queens to the same tube, following the exact same protocol. We measured one to four groups of queens per day while alternating between infected and uninfected groups (Fig S1b). Once we were done recording $CO_2$ emissions from the groups of queens for the day, we recorded a 10-min long second baseline.

The data was initially visualized and analyzed using the ExpeData software (release 1.9.13) from Sable Systems International to obtain mean values of $CO_2$ emission, humidity, and temperature over a period of stable $CO_2$ emission per experimental group. We corrected for shift between the first and secondbaseline values using ExpeData software before exporting the data as a CSV file for further analysis in R.We subtracted mean values of $CO_2$ emission of empty nests from that of the whole colonies and the brood to remove the background noise.

We compared the activity levels of whole colonies using Swarm Sight Motion Analysis software [50]. The activity data was then compiled in a CSV file to generate mean values per colony and for further analysis in R.

*Wolbachia free and genetically paired colonies*

We used six *Wolbachia*-infected and six uninfected heterogeneous stock colonies to set up reciprocal crosses (Fig S2). For reciprocal crossing, we artificially induced the production of new queens and males in six heterogenous stock colonies of known pedigree per infection status. Four weeks post this, we collected as many darkly pigmented queen and male pupae per colony as possible and stored them separately in petri dishes along with 50 workers of the same genotype. To produce genetically paired colonies, we set up a cross between 15 *Wolbachia*-infected virgin queens of genotype A and 10 uninfected virgin males of genotype B and another cross between 15 uninfected virgin queens of genotype B and 10 *Wolbachia*-infected males of genotype A (Fig. S3). Such a cross produced pairs of colonies that are genetically similar to each other but only one of them is infected with *Wolbachia* since *Wolbachia* is maternally inherited.
